# Network-based integration of epigenetic landscapes unveils molecular programs underlying human T follicular helper cell differentiation

**DOI:** 10.1101/2021.05.19.444859

**Authors:** Vinay S Mahajan, Syed A Rahman, Vinayak V Viswanadham, Grace J Yuen, Na Sun, Hamid Mattoo, Shiv S Pillai, Jishnu Das

**Affiliations:** Ragon Institute of MGH, MIT and Harvard, Cambridge, MA; Center for Systems Immunology, Departments of Immunology and Computational & Systems Biology, University of Pittsburgh, Pittsburgh, PA; Department of Biomedical Informatics, Harvard Medical School, Boston, MA

## Abstract

T follicular helper (Tfh) cells play a critical role in T-dependent humoral immune responses. While genetic programs controlling Tfh cell differentiation have been extensively studied using murine models, studies in humans have been hampered by the lack of a robust *in vitro* differentiation system for Tfh cells. We characterized epigenomic landscapes across stages of Tfh cell differentiation in a healthy human tonsil using ATAC-Seq and CUT&RUN for selected histone modifications. We combined these epigenomic datasets and integrated them with the reference human protein interactome using a novel network propagation approach. Our approach uncovered subnetworks integral to Tfh cell differentiation. These subnetworks captured known Tfh cell drivers to a greater extent than conventional gene-centric analyses would, and also revealed novel modules that may be required for Tfh cell differentiation. We find that human Tfh cell subnetworks are functionally associated with specific immune signaling cascades including cytokine receptor driven pathways. Analyses of transcriptomic data revealed that in addition to these immune pathways being significantly dysregulated in severe COVID-19, the corresponding Tfh cell subnetworks are also transcriptionally perturbed to a similar extent. This provides a molecular mechanistic basis for the previously observed impaired Tfh cell differentiation and loss of germinal centers in severe COVID-19.

## Introduction

Tfh cells are a subset of CD4^+^ T cells that help B cells enter follicles, and are critical for the development of germinal centers (GCs), somatic hypermutation and affinity maturation. Some, but not all, isotype-switching events are also mediated by Tfh cells. They are thus integral to the generation of protective humoral immune responses against pathogens. Dysregulation of Tfh cells has also been implicated in autoimmunity and cancer. Analyses of patients with inherited genetic disorders has revealed the importance of genes such as *SAP, ICOS, IL21R*, and *STAT3* in human Tfh cell differentiation and function (Crotty, 2011; Ma et al., 2012a; Schmitt et al., 2013). Although the functions of Tfh cells and their roles in working cooperatively with B cells are broadly similar in mice and humans, there are several differences in the underlying regulatory pathways, and some of these differences have already been described (Crotty, 2014; Schmitt et al., 2009, 2014). Thus, insights obtained from the study of Tfh cells and their genetic regulation using mouse models can only provide partial insights into the underlying regulatory circuits in humans.

Genetic perturbation studies in human Tfh cells have been limited by the lack of a robust *in vitro* differentiation system. In this study, we attempt to overcome this using multi-omic integration of epigenetic marks of transcriptional regulation, including those for promoters and enhancers, across the differentiation trajectory of human tonsillar Tfh cells. The multi-scale approach allows us to probe the complex underlying biology of gene regulation at multiple levels of organization. While a promoter overlaps with the transcription start site, enhancers may lie hundreds of kilobases away and influence transcription by long-range interactions. Regions of chromatin that bear a dense packing of active enhancers may be highly adept at regulating the transcription of target genes by virtue of their ability to concentrate transcription factors and transcriptional machinery in phase-separated condensates. Thus, genomic neighborhoods may have an outsized influence on enhancer function. The above factors, combined with the dynamic and cell-type-specific nature of these regulatory relationships, present a key challenge to the interpretation and combination of multi-scale epigenomic datasets, despite recent advances in techniques to map long-range enhancer-promoter interactions (Avsec et al.; Fulco et al., 2016, 2019; Nasser et al., 2021; Whalen et al., 2016; Zuin et al., 2021).

We circumvented some of these challenges using a network-based approach to integrate multi-omic epigenetic datasets. Protein interactome networks have been widely used to uncover specific molecular phenotypes underlying human genetic disorders (Vidal et al., 2011). We have demonstrated how protein networks can be combined with a range of functional genomic datasets to uncover the molecular basis of genetic disorders (Das et al., 2013, 2014; Fragoza et al., 2019; Vo et al., 2016; Wang et al., 2012; Wei et al., 2014). Others have also studied germline or somatic mutations in the contexts of protein networks, and subnetworks most affected by these perturbations (Yi et al., 2017). In cancer genomics, a range of network approaches have been used to identify driver mutations within a sea of passenger mutations (Hofree et al., 2013; Horn et al., 2018; Leiserson et al., 2015). At a conceptual level, tumor mutations and features derived from epigenomic datasets drive large-scale cell behavior in similar ways. Individual mutations or chromatin features may exert a weaker, more localized influence, but groups of mutations or epigenetic marks will exert a stronger, systemic influence by influencing multiple components of the underlying molecular network. Thus, both mutations and epigenetic features represent biological priors that are weak or uninformative when considered in isolation but are strong when analyzed together in the context of the underlying topological network structure. A network-based approach can be used to filter out isolated noisy signals and hone in on biologically significant pathways of genes that mutations or epigenomic changes would target at multiple points.

We implemented a novel network-based approach to propagate epigenomic signals derived from our ATAC-seq and CUT&RUN studies and thus uncover network modules that are likely to drive Tfh cell differentiation. Our approach is based on the premise that biologically relevant epigenetic signatures reflect the need for coordinated regulation of genes that encode interacting proteins. A similar conceptual premise has been used to identify subnetworks enriched for mutations in cancer genomics and network-based GWAS (Cowen et al., 2017; Leiserson et al., 2013, 2015; Reyna et al., 2018; Vandin et al., 2011). The underlying technique of network propagation represents a powerful way to combine a wide range of signals taking into account the structure of the underlying network. We implemented network propagation using random walk with restart, which is equivalent to an insulated heat diffusion process ((Cowen et al., 2017; Leiserson et al., 2013, 2015; Reyna et al., 2018; Vandin et al., 2011)). We used this approach to unify features derived from the different epigenomic datasets and discover networks driving Tfh cell differentiation. Our network propagation approach is different from and complementary to methods such as Taiji that prioritize individual transcription factors (TFs) (Yu et al., 2017), since those methods focus on identifying specific driver TFs from local gene regulatory networks. In contrast, we use the global reference protein interactome to uncover entire network modules, which may offer better proxies for molecular phenotypes, rather than individual TFs underlying Tfh cell differentiation.

In this study, we have established that our integrated network-based approach more completely captures known Tfh cell drivers than conventional gene-centric analyses on individual epigenomic or transcriptomic datasets. We have also uncovered novel modules that include higher-order dependencies (*i.e*., the underlying connectivity structure) not identified by genecentric approaches. We show here that the identified Tfh cell subnetworks are functionally linked to specific immune signaling pathways. These pathways are perturbed in severe COVID-19 and impinge on the corresponding Tfh cell subnetworks, dysregulating these modules to a similar extent. Our findings provide a plausible molecular mechanistic basis, centered on impaired Tfh cell differentiation, for a previously observed loss of GCs in severe COVID-19.

## Results

### Elucidating the epigenomic landscape of human Tfh cell differentiation

We systematically characterized the epigenomic landscape across the trajectory of human Tfh cell differentiation in a healthy adult human tonsil. We measured chromatin accessibility using ATAC-Seq (Buenrostro et al., 2013, 2015) and histone H3 modifications using CUT&RUN (Skene and Henikoff, 2017) with antibodies against H3K4me1, H3K4me3 and H3K27Ac. Each of these epigenomic datasets were collected at four stages of Tfh cell differentiation - naive, early pre-Tfh, late pre-Tfh and GC Tfh cells. CD4, CD45RA, CXCR5 and PD1 were used as markers to define these four subsets (Fig. 1a) as follows: CD4^+^CD45RA^+^CXCR5^-^PD1^-^ (naïve CD4+ T cells) CD4^+^CD45RA^-^CXCR5^+^PD1^-^ (early pre-Tfh cells), CD4^+^CD45RA^-^CXCR5^+^PD1^+^ (late pre-Tfh cells) and CD4^+^CD45RA-CXCR5^++^PD1^++^ (GC Tfh cells). While murine Tfh cells have been studied extensively, this resource represents a global epigenomic roadmap of human Tfh cells and offers insights derived from network biology into the regulation of human Tfh cell differentiation (Fig. 1b)

**Figure 1.**
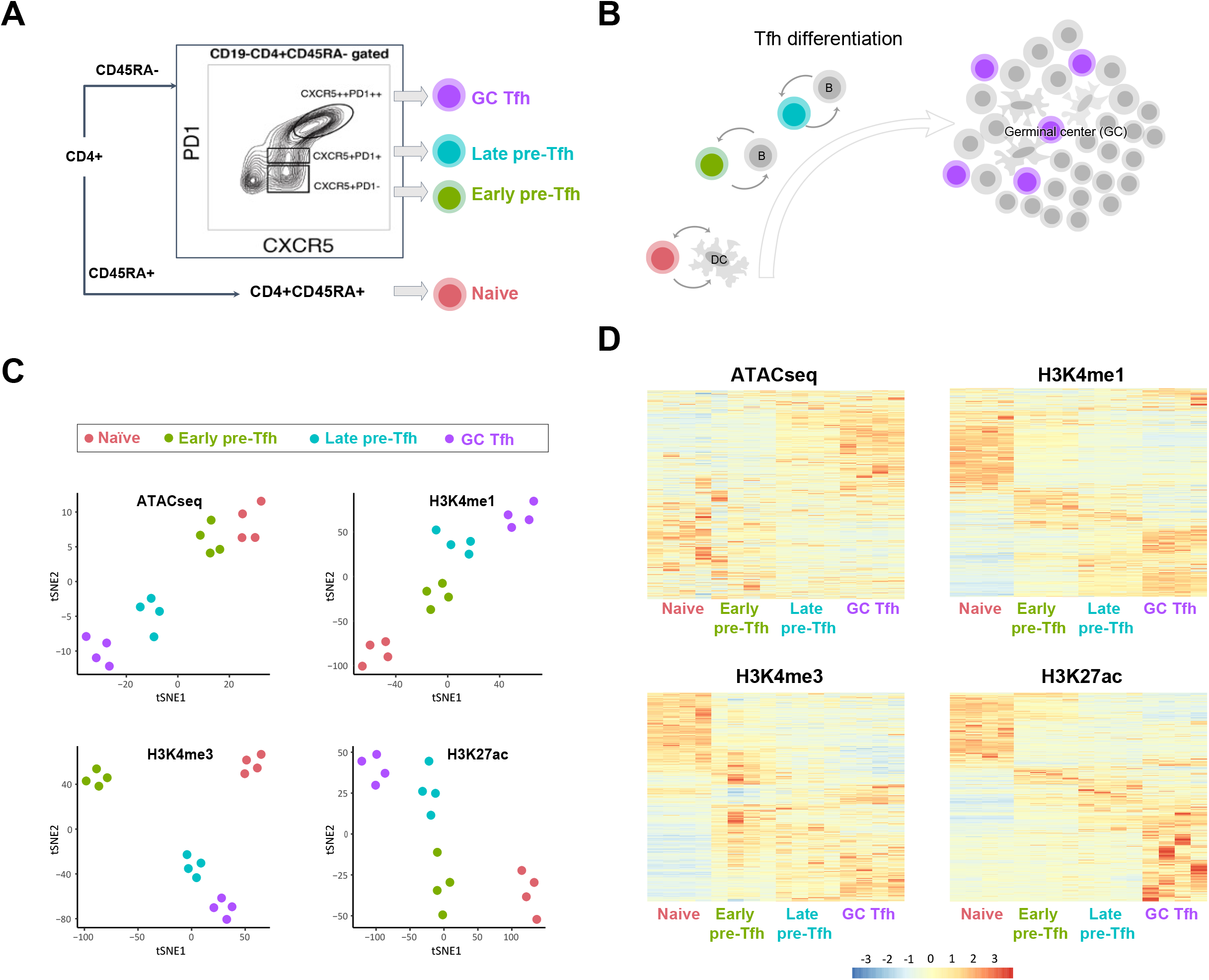
The epigenomic landscape of human Tfh cell differentiation from naive to germinal-center, and two intermediate Tfh subpopulations. A. Gating strategy to sort Tfh cells into naive, early pre-Tfh, late pre-Tfh and GC Tfh stages B. Schematic of Tfh cell differentiation and flow cytometry strategy. C. t-SNE visualizing global epigenomic profiles (ATAC-seq, H3K4me1, H3K4me3 and H3K27Ac) of samples at different stages in the Tfh cell differentiation trajectory D. Heatmap visualizing differentially regulated regions for each of the four epigenomic datasets - ATAC-seq, H3K4me1, H3K4me3 and H3K27Ac

We first analyzed each epigenomic dataset in isolation and identified significant peaks using MACS2 and organized the peaks that were less than 10 kb apart into peak clusters (Methods). For each dataset, we then visualized these peak clusters using unsupervised dimensionality reduction techniques (Fig. 1c, Fig. S1A). We observed significant stage-specific differences in low-dimensional representations of the samples for each chromatin assay type. These differences remained consistent across replicates (Fig. 1c, Fig. S1A). and suggest that these epigenomic datasets are well suited to capture the trajectory of human Tfh cell differentiation (Fig. 1c, Fig. S1A). Visualizing the differentially-marked genomic intervals for each dataset, we observed clear dynamic shifts at intervals across the four stages, with subsets corresponding to monotonically increasing or decreasing intensity in marks, or to more variegated patterns indicative of stagespecific epigenetic features (Fig. 1d). The same relationships held for bulk RNA-seq data obtained from Tfh cells at the same stages of differentiation (Fig. S1B). Together, our data captures the regulatory dynamics across Tfh cell differentiation.

### Identifying likely trajectories of Tfh cell differentiation from epigenomic data

Each of the epigenetic signals analyzed exhibited distinct trends across the four stages of Tfh cell differentiation. These differences likely reflect complementary regulatory processes underlying the differentiation of this polarized cell type (Fig. 1d). We expect that these processes all involve regulating the expression of an underlying set of components that drive the transition from naive to GC Tfh, and multi-omic integration strategies will be required to uncover these core genes. We focused on the discovery of trajectories using the epigenomic data. The transcriptomic data was used as an orthogonal dataset to validate the identified trajectories.

We first used an empirical Bayesian approach (Leng et al., 2013) to identify differentiation patterns that best fit the epigenomic data. This is an unbiased data-driven way to cluster the four states into plausible differentiation trajectories, without making *a priori* assumptions about a specific path (Fig. 2a). From the four epigenomic datasets, we recovered three differentiation patterns to which most peaks were assigned. The first pattern corresponded to a trajectory where the combined epigenomic datasets suggested there was a shared profile for naive and early pre-Tfh and a different shared profile for late pre-GC and GC Tfh cells (Fig. 2a). A second pattern corresponded to distinct profiles for all four stages (Fig. 2a). The third pattern corresponded to naive CD4+ T cells versus the other three stages. We focused on the first two, as they relate to likely trajectories of Tfh cell differentiation while the third is likely to reflect epigenomic differences between naive and non-naive cells, which was not the focus of this study.

**Figure 2.**
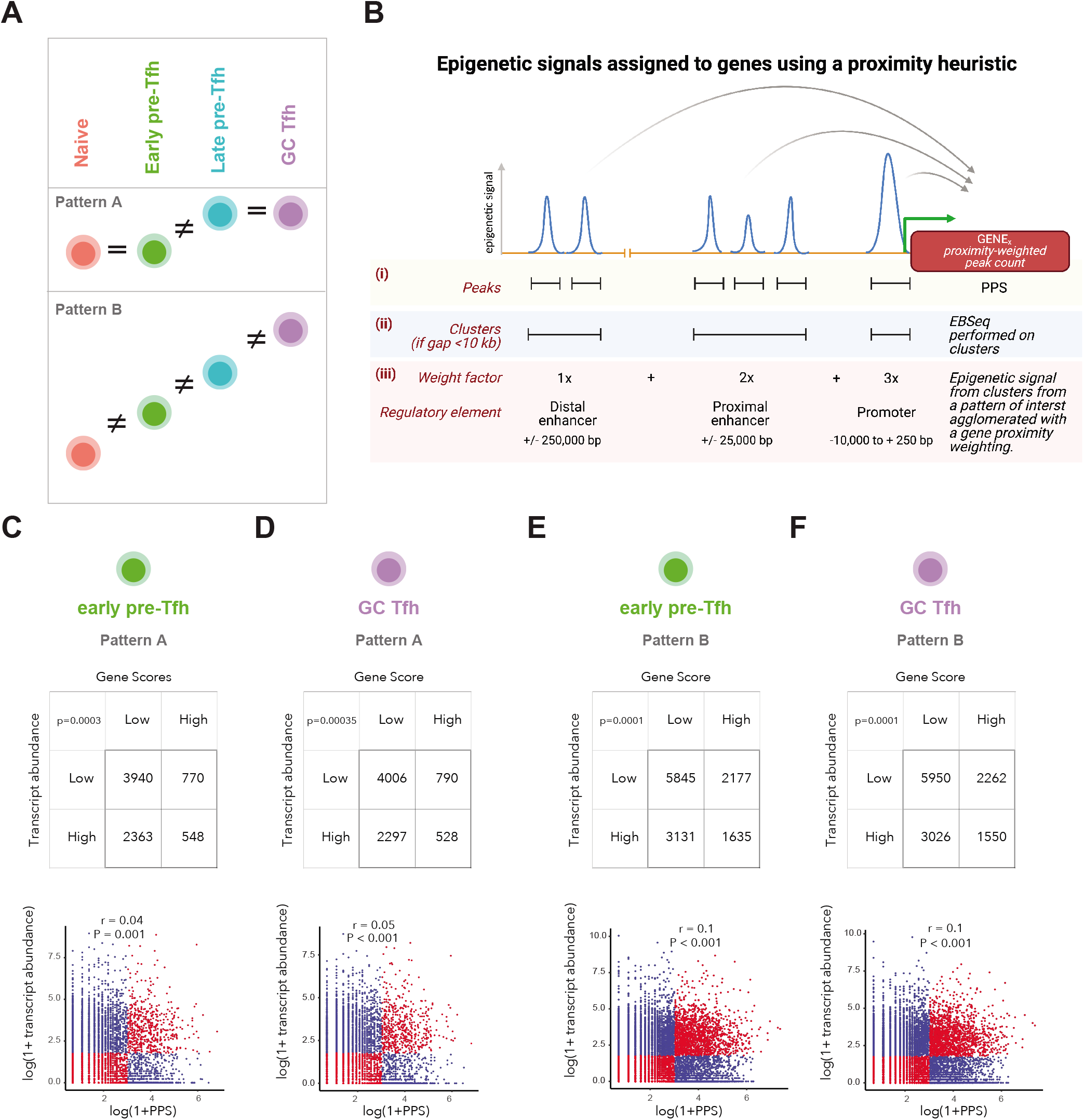
Calculating gene-centric peak proximity scores (PPS’s) to integrate epigenomic data from differentiating Tfh cells. A. Patterns of Tfh cell differentiation identified using EBSeq. B. Schematic of a proximity-weighted count heuristic to calculate peak proximity scores. C. Relationship between PPS and transcript abundances at the early pre-Tfh stage for pattern A of Tfh cell differentiation (PPS threshold 20, transcript abundance threshold 5) D. Relationship between PPS and transcript abundances at the GC pre-Tfh stage for pattern A of Tfh cell differentiation (PPS threshold 20, transcript abundance threshold 5) D. Relationship between PPS and transcript abundances at the early pre-Tfh stage for pattern B of Tfh cell differentiation (PPS threshold 20, transcript abundance threshold 5) F. Relationship between PPS and transcript abundances at the GC pre-Tfh stage for patternB of Tfh cell differentiation (PPS threshold 20, transcript abundance threshold 5)

Next, for each differentiation pattern, peaks from each dataset were aggregated into gene-centric scores using a proximity-weighted count heuristic (Fig. 2b) based on first principles underlying the organization of promoters and enhancers. Gene-centric scores from each dataset were then combined by weighting the datasets equally, as we did not have any *a priori* information regarding relative information context across the datasets. We refer to each gene’s score as its “peak proximity score” (PPS), which we can calculate for each possible differentiation pattern and use as a surrogate for gene regulation. We then sought to evaluate whether each gene’s PPS tracked with corresponding transcript abundances for Tfh cells prior to entering the GC (at the early pre-GC stage) or at the GC Tfh stage. For both differentiation patterns, we observed that there was a weak correspondence between the scores and transcript abundances at each stage (Figs. 2c–2f). Gene-centric scores stratified into high/low had a significant correspondence with transcript abundances stratified into high/low (Figs. 2c–2f). This relationship remained consistent at different thresholds to dichotomize gene-centric scores and transcript abundances (Figs S2A-S2F). However, the Spearman correlations between the gene-centric scores and transcript abundances (Figs 2c–2f) were weak. These suggest that although individual gene-centric scores capture specific aspects of gene regulation, in isolation they serve only as incomplete priors for uncovering complex regulatory relationships.

### Network-based integration of epigenomic datasets uncovers coordinated modules driving Tfh cell differentiation along identified trajectories

Our findings motivated us to examine whether these individual gene-centric scores could be combined into a strong biological signal by accounting for additive effects and dependencies. Specifically, we incorporated higher-order relationships between proteins encoded by these genes using the high-quality (i.e., each edge is validated experimentally using multiple independent assays) reference human protein interactome network (Cusick et al., 2009; Das and Yu, 2012).

We ran network propagation through HotNet2 using the combined PPS’s for each protein’s gene as seed values (Figs. 3a, 3b). HotNet2 helps us incorporate the topological structure of the underlying protein interaction network as an informative prior on how proteins are functionally related to one another. The use of network propagation is motivated by the hypothesis that proteins encoded by relevant genes relevant to Tfh cell differentiation are more likely to interact with one another and make up cohesive network submodules (Fig. 3b). Further, our approach can help filter out genes that may not play a role in Tfh cell differentiation but nevertheless show similar epigenetic patterns to relevant genes. Thus, network propagation constitutes the final step in a three-step pipeline: identification of the most likely patterns of Tfh cell differentiation, generation of gene-centric scores using a proximity-weighted count heuristic (i.e. calculation of PPS’s), and network propagation using these scores as seeds.

**Figure 3.**
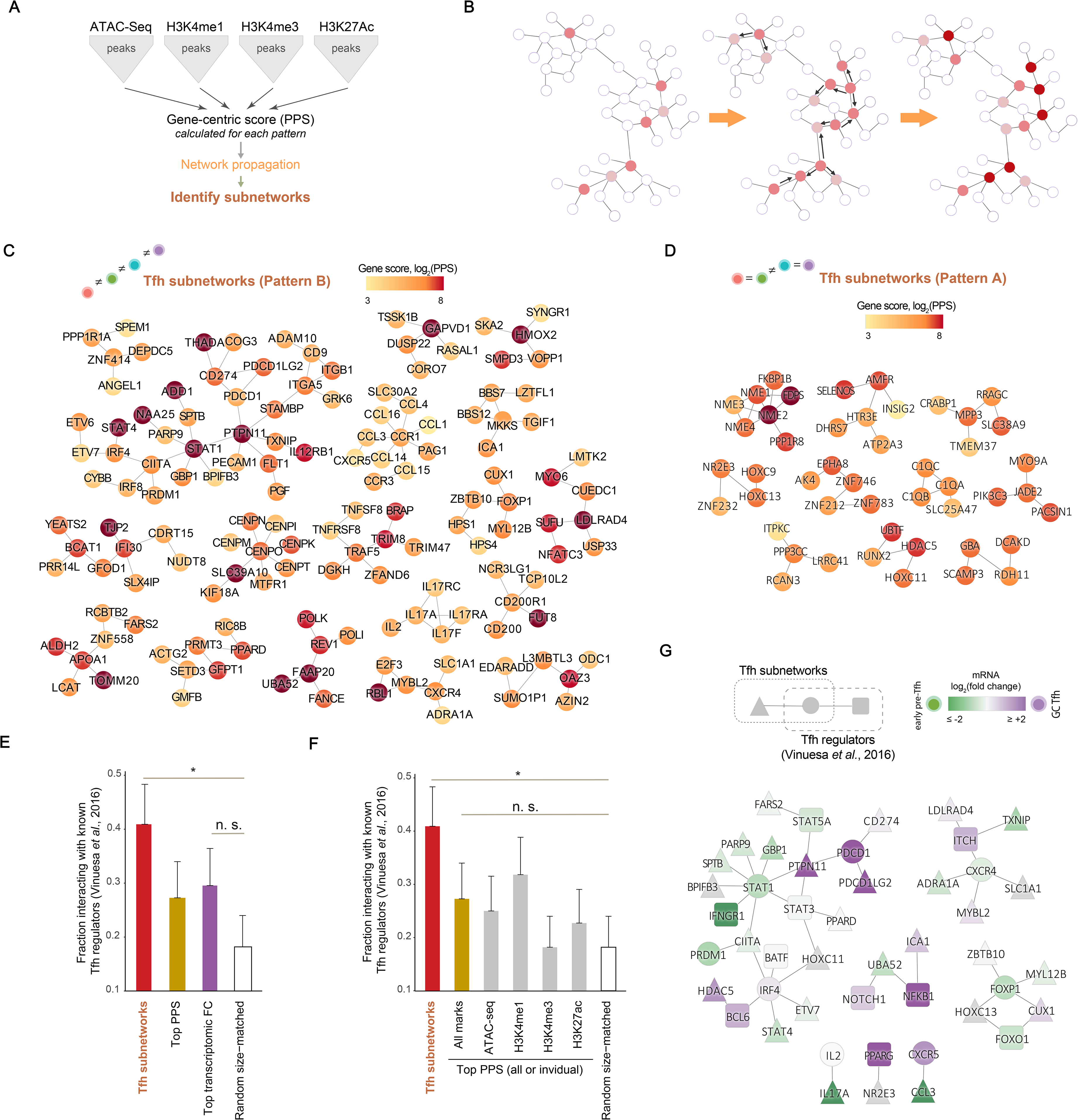
Network-based integration of epigenomic datasets uncovers coordinated subnetworks driving Tfh cell differentiation along identified trajectories. A. Datasets used in the integrative network analyses. B. Schematic illustrating network propagation C. Tfh cell subnetworks corresponding to differentiation pattern B D. Tfh cell subnetworks corresponding to differentiation pattern A E & F. Fraction of proteins in different gene sets that functionally overlap with known drivers of Tfh cell differentiation G. Functional overlap between Tfh cell subnetworks and known drivers of Tfh cell differentiation (colored by logFC along the differentiation trajectory)

By propagating these scores, we converged on a set of coherent modules for each of the two patterns of interest underlying the Tfh cell differentiation trajectory (Figs 3c, 3d Methods). The statistical significance of the identified subnetworks was assessed using network permutations as described in HotNet2. The identified Tfh cell subnetworks are both significant when compared to controls generated using permuted networks and stable across technical replicate network propagation runs (Methods). Next, we validated the biological significance of these subnetworks by calculating whether our subnetwork shares or directly interacts with proteins from a set of experimentally-validated drivers of Tfh cell differentiation (Vinuesa et al., 2016). We found that the Tfh cell subnetworks had significantly higher overlap with experimentally-validated regulators of Tfh cell differentiation compared to a random size-matched control of network genes (Fig. 3e). More interestingly, a size-matched set of prioritized genes chosen based on top PPS (combined across the 4 epigenomic datasets) prior to network propagation had a lower overlap with known drivers of Tfh cell differentiation, compared to the Tfh cell subnetworks (Fig. 3e). A similar trend was observed with a size-matched set of genes prioritized based on fold changes from our transcriptomic dataset (Fig. 3e). The trend remained unchanged if we utilized a size-matched set of prioritized genes chosen based on the top PPSs from each of the 4 individual epigenomic datasets (Fig. 3f). Our results show that while there is some signal (numerically higher than a random size matched control) in gene-centric PPS’s derived from epigenomic and transcriptomic datasets, a network-based approach that combines these PPS’s performs much better at recapitulating known drivers of Tfh cell differentiation that gene-centric PPS’s derived from individual or combined epigenomic or transcriptomic datasets (Figs. 3e, 3f).

Next we examined if we recovered known trajectories beyond simply overlapping with known drivers. We used fold changes across the differentiation trajectory, obtained from the RNA-seq data to visualize the obtained trajectories. We recovered several well-known components including an antagonistic relationship between BCL6 and PRDM1 (BLIMP1), a known master regulator circuit governing Tfh cell differentiation (Fig. 3g, Fig. S3) (Johnston et al., 2009; Vinuesa et al., 2016). We also recovered circuits involving IRF4, STAT1, PDCD1 (PD1) and CXCR5 (Fig. 3g, Fig. S3). Importantly, in addition to recovering known interactions amongst genes in Tfh cells, we identified several novel relationships. Of specific interest was an intriguing connection between the BCL6/PRDM1/IRF4/STAT1 module and the PD1 subnetwork; this connection involved PTPN11 (SHP2), which is known to be activated by PD1 (Marasco et al., 2020). Our results suggest that a previously characterized crosstalk between the BCL6/PRDM1/IRF4/STAT1 and PD1 modules mediated by PTPN11 regulates Tfh cell differentiation.

### Tfh cell subnetworks are functionally associated with specific immune signaling pathways

The identified Tfh cell subnetworks capture known regulators of Tfh cell differentiation better than individual epigenomic or transcriptomic datasets and uncover cell differentiation patterns not reflected at the level of individual genes (Fig. S4). Next, we sought to evaluate whether there are significant functional overlaps (i.e., shared or directly interact with) between these proteins in these subnetworks and Hallmark, KEGG and PID pathways (Subramanian et al., 2005). For each of these pathway databases, we calculated the functional overlap between the Tfh cell subnetworks and all pathways in the corresponding database. We assessed the significance of these overlaps using a size-matched random control - this represents the baseline overlap between a random size-matched set of proteins from the network and these pathways. We visualized these using volcano plots, where each dot represents the extent/effect size (quantified using log odds ratio on the X axis) and significance (quantified using -log P on the Y axis) of the functional overlap between the Tfh cell subnetworks and a particular pathway in the database (Methods).

Interestingly, of the 50 Hallmark pathways, we saw that only a single pathway, IL6-JAK-STAT signaling, significantly functionally overlapped with the Tfh cell subnetworks (Figs 4a, 4b). The enrichment results overall suggest that the Tfh cell subnetworks have specific functional associations, confirming their specificity. With KEGG pathways, we also observed the functional overlap with IL6-STAT signaling (Figs. 4c, 4d), demonstrating that our analyses robustly capture the same IL6 pathway as defined in two different databases. We also identified a significant functional overlap with the KEGG cytokine receptor signaling pathway (Figs 4c, 4d). The significant overlaps were also very specific (2 out of 186 KEGG pathways). Finally, we observed a significant functional overlap with 2 of 196 PID pathways: IL12-STAT4 and integrin signaling (Figs 4e, 4f). Overall, our analyses hone in on very specific immune signaling pathways that are functionally associated with the identified Tfh cell subnetworks, providing context to processes with which they are associated.

**Figure 4.**
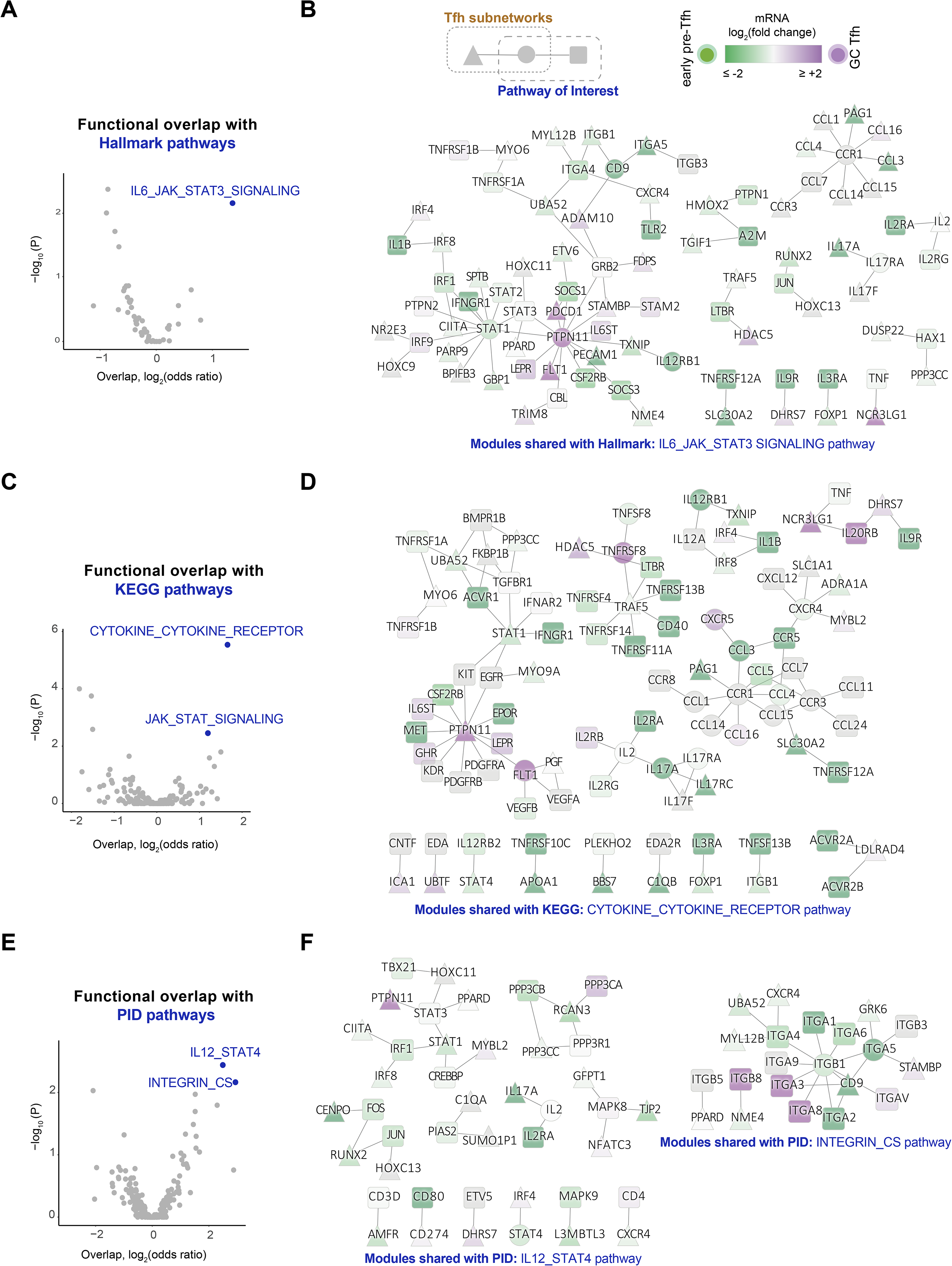
Tfh cell subnetworks identified by network-based integration functionally overlap with specific immune and cytokine signaling pathways. A. Volcano plot illustrating effect sizes and significance of functional overlaps between Tfh cell subnetworks and Hallmark pathways. B. Functional overlap between Tfh cell subnetworks and the Hallmark IL6_JAK_STAT pathway (colored by log FC along the differentiation trajectory) C. Volcano plot illustrating effect sizes and significance of functional overlaps between Tfh cell subnetworks and KEGG pathways. D. Functional overlap between Tfh cell subnetworks and the KEGG CYTOKINE_CYTOKINE_RECEPTOR pathway (colored by log FC along the differentiation trajectory) E. Volcano plot illustrating effect sizes and significance of functional overlaps between Tfh cell subnetworks and PID pathways. F. Functional overlap between Tfh cell subnetworks and the PID IL12_JAK_STAT & INTEGRIN_CS pathways (colored by log FC along the differentiation trajectory)

Some of the identified associations are well known, with evidence in mechanistic studies including *in vivo* murine models. For example, the IL6 association of the Tfh cell subnetworks is well known as circulating plasmablasts are known to induce cell Tfh cell differentiation by IL6 production (Chavele et al., 2015). And consistent with our findings, integrins have been shown to have an important role in sustained Tfh support of the GC response, which in turn is important for the generation of long-lived plasma cells (Schrock et al., 2019). However, some of the other identified associations are counterintuitive and novel. The identified overlap between the Tfh cell subnetworks and IL12/STAT signaling suggests that this association negatively regulates Tfh cell differentiation (Fig. 4e). While this would appear to be counter-intuitive in light of previous reports, recently, in a mouse model, IL12 has been shown to have a key role in blocking Tfh cell differentiation and contributing to GC suppression (Elsner and Shlomchik, 2019). Furthermore, we uncover a novel role of PTPN11 in Tfh cell differentiation as it emerges as a key hub in several subnetworks (Figs. 4b, 4d, 4f). The Tfh cell subnetworks identified in this study provide a global systems perspective to the roles of signaling cascades, some of which have been individually elucidated earlier, and others that are novel, in Tfh cell differentiation.

### Pathways dysregulated in severe COVID-19 impinge on Tfh cell subnetworks

To further understand the implications of the functional overlaps, we sought to evaluate whether these molecular phenotypes could explain organism-level phenotypes involving dysregulation of Tfh cell differentiation. A striking example of such a phenotype was reported recently in severe COVID-19, where Tfh cells and GCs were observed to be completely lost in severe COVID-19 (Kaneko et al., 2020). We hypothesized that the loss of Tfh cells in severe COVID-19 could be linked to transcriptional dysregulation of the immune signaling pathways functionally associated with the significant Tfh cell subnetworks (Fig. 5a). The dysregulated immune signaling pathways would then impinge on the Tfh cell subnetworks, blocking the differentiation of Tfh cells into the most differentiated GC Tfh form (Fig. 5a).

**Figure 5.**
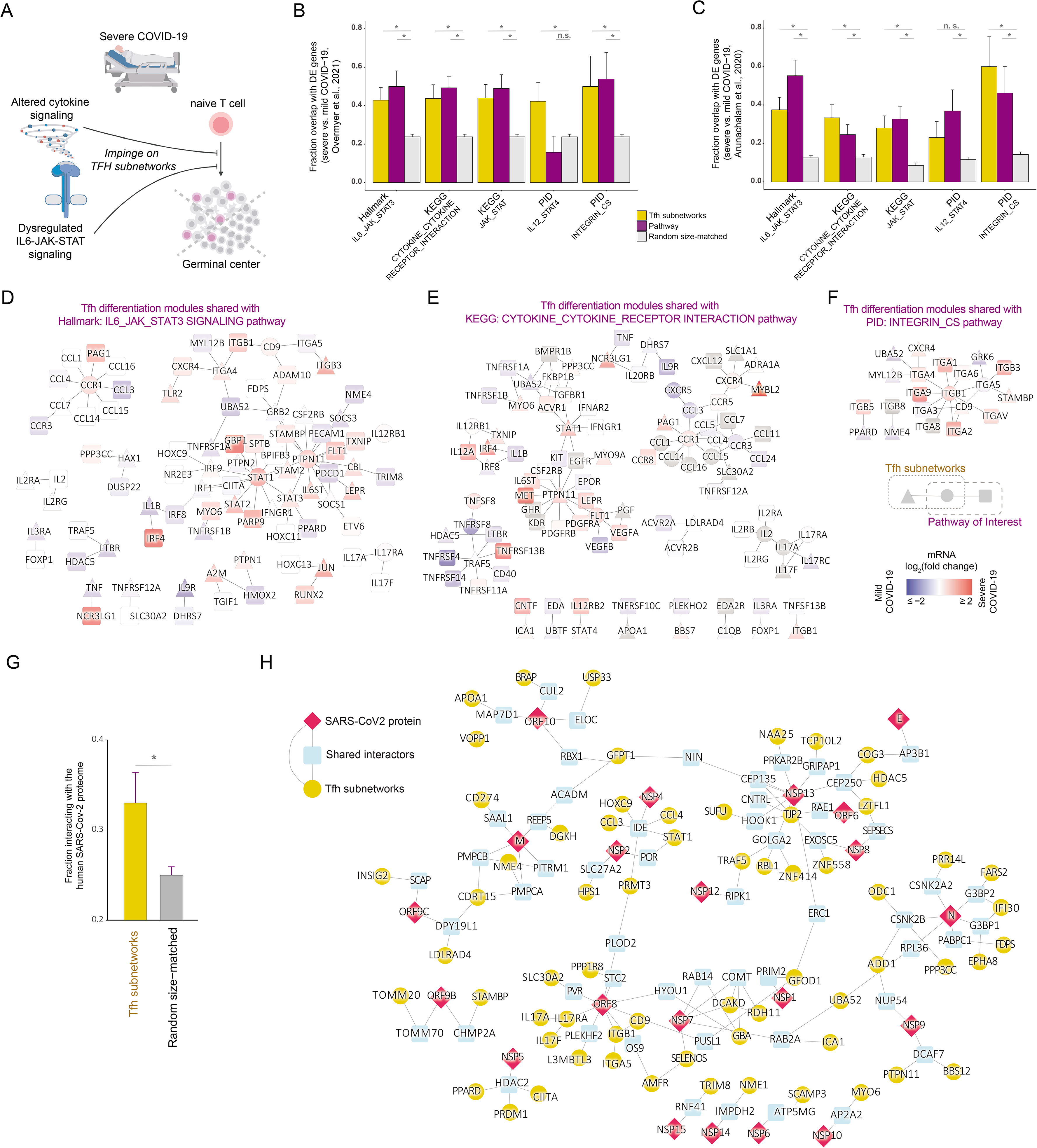
Pathways dysregulated in severe COVID-19 impinge on the Tfh cell subnetworks. A. Schematic of how pathways dysregulated in severe COVID-19 impinge on the Tfh cell subnetworks B. Fraction of significant DEGs from Overmyer et al (Overmyer et al., 2021) between severe and mild COVID-19 in gene sets of interest C. Fraction of significant DEGs from Arunachalam et al (Arunachalam et al., 2020) between severe and mild COVID-19 in gene sets of interest D. IL6_JAK_STAT pathway (Hallmark) impinging on the Tfh cell subnetworks (colored by log FC between severe and mild COVID-19) E. Cytokine_cytokine_receptor pathway (KEGG) impinging on the Tfh cell subnetworks (colored by log FC between severe and mild COVID-19) F. Integrin_CS pathway (PID) impinging on the Tfh cell subnetworks (colored by log FC between severe and mild COVID-19) G. Fraction of proteins in the Tfh cell subnetworks that functionally overlap with the SARS-CoV2-human interactome H. Functional overlap between the SARS-CoV2-human interactome and the Tfh cell subnetworks

While the Tfh cell subnetworks themselves are expressed in human Tfh cells (Fig. S5), we used publicly available PBMC transcriptomic datasets from COVID-19 patients to test our hypothesis, as the crosstalk between dysregulated immune signaling and blocked cell Tfh cell differentiation could involve both cell-intrinsic (i.e., within Tfh cells) and cell-extrinsic (i.e., involving cross-talk between Tfh cells and other immune cells) mechanisms. We used two orthogonal PBMC transcriptomic datasets from different geographical locations, both of which had identified significant DEGs between severe and mild COVID-19 patients (Arunachalam et al., 2020; Overmyer et al., 2021). For both datasets, we found that DEGs were significantly overrepresented (compared to a size-matched set of random genes) in the immune signaling pathways (IL6-JAK-STAT, IL12-STAT4, cytokine receptor and integrin signaling) functionally associated with the Tfh cell subnetworks (Figs. 5b, 5c). This was unsurprising, as multiple reports have regulated dysregulation of these pathways in severe COVID-19 (Lei et al., 2020). However, more surprisingly, we found that the DEGs were also significantly over-represented in components of the Tfh cell subnetworks that functionally overlap with each of these immune pathways (Figs. 5b, 5c). Our findings show that DEGs between severe and mild COVID-19 patients are over-represented both in specific immune signaling pathways and the corresponding functionally-associated Tfh cell subnetworks. This suggests they are both similarly transcriptionally dysregulated in severe COVID-19 (Figs 5b, 5c). Interestingly, the DEGs are spread through the entire modules, rather than being limited to individual genes (Figs. 5d–5f). This suggests that the network approach helps us identify specific functional processes (modules) involving cross-talk between immune signaling and Tfh cell differentiation that are transcriptionally dysregulated in COVID-19 (Figs. 5d–5f). Further, while the identified Tfh cell subnetworks are expressed in human Tfh cells (Fig. S4), the identification of transcriptionally perturbed modules from a PBMC-based transcriptomic dataset allows us to identify both cell-intrinsic and cell-extrinsic signaling cascades altered in severe COVID-19. Overall, our findings support the prior hypothesis that the loss of GCs and Tfh cells in severe COVID-19 are linked to aberrant Tfh cell differentiation (Kaneko et al., 2020). Further, we provide the first molecular mechanistic evidence in support of this hypothesis via the identification of specific immune signaling pathways that impinge on the Tfh cell subnetworks, blocking the Tfh cell state in severe COVID-19.

Finally, we examined the functional overlaps between the Tfh cell subnetworks and the SARS-Cov2 host-viral protein interactome (Gordon et al., 2020). We uncovered a significant functional overlap driven by proteins in the Tfh cell subnetworks that interact with the human interactors of the viral proteins, but not the viral proteins themselves (Figs. 5g, 5h). It is not known how exactly the Tfh cell subnetworks interact with viral proteins, but it raises the intriguing possibility that the viral proteome may directly influence Tfh cell differentiation via its interacting partners, including proteins expressed in other cell types.

## Discussion

In this study, we applied a novel network propagation approach to integrate multi-omic datasets that represent gene regulation at different levels and uncover subnetworks involved in human Tfh cell differentiation. Specifically, we adapted the HotNet2 algorithm to identify protein network modules enriched for signals derived from ATAC-seq and CUT&RUN datasets obtained from Tfh cells at four stages of differentiation within a healthy human subject’s tonsil. By framing the search for drivers as a search for identifying protein subnetworks with concentrated epigenomic signals, our approach recovered known regulators of Tfh cell differentiation, and we have implicated several novel candidate genes involved in Tfh cell differentiation. Within the identified Tfh cell subnetworks, we found functional associations between Tfh cell differentiation and specific immune signaling pathways including IL6, IL12, and JAK-STAT. While some of the identified associations such as those with IL6 and integrin signaling are well known, others such as the role of IL12 as a negative regulator of Tfh cell differentiation or PTPN11 as a key hub (connecting the BCL6/PRDM1/IRF4/STAT1 and PD1 modules) are novel.

We find that these pathways are significantly altered in severe COVID-19 and perturb interactions amongst proteins within Tfh cell subnetworks, likely impairing Tfh cell differentiation. We have previously described the striking loss of GCs in lymph nodes and spleens and depletion of Bcl6+ B cells in COVID-19 patients (Kaneko et al., 2020). This finding led us to hypothesize that a specific block in GC type Bcl6+ T follicular helper cell differentiation may explain the loss of GCs and the accumulation of non-GC-derived activated B cells, and which may provide an underlying basis for the lower quality and lack of durability of humoral immune responses observed during natural infection with SARS-CoV-2. Here, we provide a plausible molecular mechanistic basis for the blocked Tfh cell differentiation. Our data leads us to hypothesize that this block arises from transcriptomic dysregulation in the identified immune signaling pathways impinging on the corresponding Tfh cell subnetworks. From our analysis, IL6 and IL12 signaling components appear at the heart of subnetworks underlying Tfh cell differentiation. Two recent separate studies suggest that increased IL6 levels in the plasma are a biomarker of severe COVID-19 (Filbin et al., 2020; Laing et al., 2020). While IL6 is a broadly relevant pro-inflammatory cytokine, the increased levels of IL6 from severe COVID-19 are likely to play a role in altering Tfh cell development. Our analysis suggests that IL6 and IL12 signaling work through a complex network of interactions to regulate Tfh formation, with our specific results indicating that the activity of centrally connected components in our subnetworks such as STAT1 may either promote or inhibit Tfh cell differentiation in different contexts. Specifically, while the expression patterns of centrally connected nodes suggest that they are upregulated during the progression to Tfh, these nodes and their interactors are often overexpressed in severe COVID-19. Coupled with our observations about the loss of GCs in severe COVID-19, our results suggest that aberrant activity of these components contribute to Tfh loss. Thus, Tfh cell differentiation pathways are uniquely sensitive to the gene expression perturbations in gene expression observed in severe COVID-19 relative to mild disease, thereby providing additional evidence that the loss of GCs in severe COVID-19 is the likely direct or indirect result of impaired Tfh cell differentiation. Our work also further supports the hypothesis that IL6 plays a significant role in the prognosis of severe versus mild COVID-19.

A block in GC differentiation has also been observed in a number of murine infection models, including *Salmonella enterica* serovar *Typhimurium* (Elsner and Shlomchik, Cell Reports 2019), *Ehrlichia muris* (Popescu et al., 2019), and *Plasmodium berghei* (Ryg-Cornejo, Cell Reports 2016). Furthermore, in a mouse model of chronic LCMV infection following DC vaccine-elicited CD4 T cells, the immunized mice exhibit loss of GCs in the setting of severe lung inflammation and a virally induced cytokine storm characterized by both lymphopenia, a dramatic loss of GCs and Tfhs, resembling severe COVID-19 infection (Penaloza-McMaster, Science 2015). However, the specific molecular factors leading to the loss of GCs were not explored in this study. While the loss of GCs and Tfhs in the *E. muris* murine infection model could be reversed by TNFα blockade, dual blockade of TNFα and IFNγ during *P. berghei* infection was required for optimal reversal of GC loss. Studies of Salmonella enterica serovar Typhimurium infection in mice implicated the role of IL-12 production in contributing to GC suppression via suppression of Tfh cell differentiation. As all of these murine infection studies are characterized by intense type 1 immune responses, our model supports the conclusion that type 1 immune responses involving IL-12 and STAT signaling inhibit Tfh differentiation and reveals the underlying protein interaction network that mediates this effect.

In summary, we have identified protein subnetworks that significantly contribute to Tfh cell differentiation. Our approach and findings are complementary to current analytic techniques that focus on the discovery of specific transcription factors (TFs) driving such cellular differentiation processes. While prioritized individual TFs help elucidate specific regulatory processes (Yu et al., 2017), our network approach uncovers both the roles of TFs and their protein interactors, thus capturing biological control at the regulatory and the protein network levels. More broadly, analyzing Tfh cell dynamics demonstrates that network propagation represents a flexible and broadly useful approach for studying biological processes that require the integration of disparate epigenomic datasets. This network analysis workflow will be a critical tool to identify multi-scale layers of control into gene regulation.

## Methods

### Flow cytometry and sorting

600 million cells from human tonsils were resuspended in Miltenyi magnetic-activated cell sorting (MACS) buffer and stained with biotinylated anti-human CD19 (clone: HIB19; BioLegend, #555411) on ice for 30 min. Cells were washed once in MACS buffer and incubated with antibiotin microbeads (Miltenyi Biotec) for 30 minutes on ice. Cells were then loaded on Miltenyi LS columns at 300 million cells per column, and the flow-through was collected as the B cell–depleted fraction. These cells were spun down and subjected to Fc block (Human BD Fc Block BD Biosciences, #564219), washed with 1% BSA in PBS, and stained with LIVE/DEAD Fixable Blue Dead Cell Stain (ThermoFisher Scientific, #L23105), before they were surface stained with antibodies against CD3 (clone: SK7; BioLegend, #344817), CD4 (clone: OKT4; BioLegend, #317420), CD45RA (clone: H100; BioLegend, #304122), CXCR5 (clone: J252D4; BioLegend, #356920), and PD1 (clone: EH12.2H7; BioLegend, #329924). The following populations were sorted using FACSAria2 (Becton Dickinson), as shown in the gating strategy present in Fig. 1A: naive CD4 T cells (CD4+CD45RA+CXCR5-PD1-), (ii) early pre-Tfh (CD4+CD45RA-CXCR5+PD1-), (iii) late pre-Tfh (CD4+CD45RA-CXCR5+PD1+), and (iv) GC Tfh cells (CD4+CD45RA-CXCR5++PD1++). All antibodies were used at a dilution of 1:40.

### RNA-seq and analysis

Total RNA was isolated from the FACS-sorted healthy tonsillar cells using the RNeasy plus Micro Kit, according to the manufacturer’s instructions (Qiagen). RNA-sequencing libraries were prepared as previously described (Picelli et al, 2013). Briefly, whole transcriptome amplification and tagmentation-based library preparation were performed using the SMART-Seq2 protocol. Libraries were assessed for quality using High Sensitivity DNA chips on the Agilent Bioanalyzer, quantified using Qubit fluorometer (Thermo Fisher Scientific), as well as KAPA Library Quantification kit (KAPA Biosystems), followed by 35-bp paired-end sequencing on a NextSeq 500 instrument (Illumina). 5–10 million reads were obtained from each sample and aligned to the University of California, Santa Cruz hg38 transcriptome. Gene expression was calculated using RSEM as previously described (Li & Dewey, 2011), with alignments to ENSEMBL hg38, and differential expression analysis was conducted using EBSeq, with differentially-expressed genes determined within each pattern by selecting genes with a PPDE (posterior probability of differential expression) of at least 0.95.

### CUT&RUN

CUT&RUN libraries were prepared using rabbit monoclonal antibodies against H3K4me1, H3K4me3 and H3K27Ac (Cell Signaling Technology catalog numbers 5326, 9751, and 8173, respectively) as previously described. The protein-A Mnase fusion protein was a gift from Steven Henikoff.

### ATAC-seq

ATAC-Seq libraries were prepared as originally described (Buenrostro et al., 2013). Fifty thousand freshly sorted cells of each population were pelleted and washed with 50 μL chilled 1× PBS, and with 50 μL lysis buffer (10 mM Tris-HCl pH 7.4, 10 mM NaCl, 3 mM MgCl2, 0.1% IGEPAL CA-630). The nuclei were pelleted in a cold micro-centrifuge at 550×g for 10 min, and resuspended in a 50μL transposition reaction with 2.5μL Tn5 transposase (FC-121-1030; Illumina) to tagment open chromatin. The reaction was incubated at 37°C in a Thermomixer (Eppendorf) at 300 rpm for 30 min. Tagmented DNA was purified using a QIAGEN MinElute kit and amplified with 7 or 11 cycles of PCR, based on the results of a test qPCR. ATAC-Seq libraries were then purified using a QIAGEN PCR cleanup kit and quantified using KAPA library quantification kit (KAPA Biosystems, Roche) and sequenced on the Nextseq 550 platform.

### EBSeq pattern analyses

Peaks were called with MACS2. Peaks within 10 kb were clustered into peak clusters. Overlapping peak clusters across all stages of differentiation were merged and the epigenetic signal within this interval for each sample was used as input for EBSeq analysis (Leng et al., 2013). Differential signal in epigenomic and transcriptomic datasets was quantified using EBSeq using the standard procedure in the EBSeq manual. For analysis of peak clusters by patterns of interest, those with a PPDE (posterior probability of differential expression) less than 0.95 for the most likely pattern were filtered.

### Calculation of a gene-centric peak proximity scores

For a biological (EBSeq) pattern of interest, peak proximity scores (PPS’s) for a gene (“genecentric PPS”) were calculated for a biological pattern of interest by calculating a weighted sum of the number of peak clusters from the pattern in the gene’s vicinity using distance-based weights to reflect the varying contributions of promoter (± 10 kb) [3x weight], proximal enhancer (± 25 kb) [2x weight] and distal enhancers (± 250 kb) [1x weight] to gene regulation.

Gene-centric PPS scores were first calculated individually for each epigenomic dataset. These were then combined into a final PPS score using equal weights for the different datasets.

### Network propagation using random walk with restart

For each pattern, we used network propagation to integrate the gene-centric scores. We used the random walk with restart algorithm (equivalent to an insulated heat diffusion process) as implemented in HotNet2 (Leiserson et al., 2015) to identify high scoring sub-networks. Relevant HotNet2 code is available at https://github.com/raphael-group/hotnet2. We ran network propagation on the union of the high-quality reference human binary and co-complex protein interactomes (Das and Yu, 2012). To generate the relevant files needed to run the random walk with restart (influence matrix of the input network, permuted networks, and their corresponding influence matrices), we used the corresponding HotNet2 scripts (“makeNetworkFiles.py” function and kept the files in HDF5 format). We generated 100 network permutations and 100 heat permutations. We then ran network propagation with the default HotNet2 parameters (beta = 0.4 and 4 different deltas generated by HotNet2). We defined significant modules for each run using *P* < 0.1. We performed 5 independent runs of network propagation, and only retained significant subnetworks that showed up in all of the 5 independent runs or replicates. We termed the significant subnetworks as the Tfh cell subnetworks.

### Estimation of functional overlap between Tfh cell subnetworks and pathway databases

We examined the functional overlap (where functional overlap was defined as either shared or directly interacting with) between the Tfh cell subnetworks and known drivers of Tfh cell differentiation. We used top-scoring genes based on the individual gene-centric scores (without network propagation) and a size-matched control of random network genes for comparison. P values were calculated using a binomial proportions test. We colored these overlapping modules based on the log fold changes across the trajectories, where fold change was defined as *log*2(*S*4/*S*2), where *S*4, *S*2 refer to the expressions for stage 4 (GC) and stage 2 (early pre-GC) respectively. We also calculated the functional overlap (as defined above) between the Tfh cell subnetworks and all pathways in 3 different pathway databases - KEGG, Hallmark and PID. We calculated the significance of these overlaps based on a random size-matched set of network genes. Significant functional overlaps were defined using an odds ratio (odds of overlap of the Tfh cell subnetworks with a pathway/odds of overlap of a random size-matched set with that pathway) threshold of >2 and an FDR threshold of 0.05 (P value calculated using a binomial proportions test and adjusted for multiple comparisons using Benjamini-Hochberg multiple-testing correction). These modules were colored as described earlier (using the log fold changes as described above).

### Analysis of transcriptomic dysregulation of the Tfh cell subnetworks and relevant pathways in COVID-19

We used two different publicly available transcriptomic datasets from COVID-19 patients across the severity spectrum. Overmyer et al identified differentially regulated genes (DEGs) between mild and severe COVID-19 patients (Overmyer et al., 2021). We used this study to compute the fraction of genes in the Tfh cell subnetworks and the relevant pathways of interest that were significantly dysregulated in COVID-19 (i.e., DEGs in the Overmyer et al study). We used a size-matched set of network genes as a random control. We used a binomial proportions test to calculate the statistical significance of the comparisons. To further validate that the identified trends were not specific to a cohort or dataset, we used another orthogonal cohort from Arunachalam et al to perform a similar analysis (Arunachalam et al., 2020). We also visualized the corresponding modules based on these fold changes.

### Assessment of links between the Tfh cell subnetworks and the host-viral interactome

We investigated links between the Tfh cell subnetworks and the SARS-Cov2-human interactome (Gordon et al., 2020) by computing the number of proteins in the Tfh cell subnetworks that were shared or directly interacting with the SARS-CoV2-human interactome (i.e., directly interacting with a viral protein or directly interacting with a human interactor of a viral protein). We used a size-matched set of network genes as a random control. We used a binomial proportions test to calculate the statistical significance of the comparisons.

## Supplementary Figure Legends

**Figure S1 - Epigenomic and transcriptomic landscapes of human Tfh cell differentiation**

A. PCA visualizing global epigenomic profiles (ATAC-seq, H3K4me1, H3K4me3 and H3K27Ac) of samples at different stages in the Tfh cell differentiation trajectory

B. Heatmap visualization of differentially expressed genes for the bulk RNAseq dataset

**Figure S2 – Relationship of PPS’s and transcript abundances at different thresholds**

A. At the early pre-Tfh and GC-Tfh stages for pattern A of Tfh cell differentiation (PPS threshold 50, transcript abundance threshold 10)

B. At the early pre-Tfh and GC-Tfh stages for pattern A of Tfh cell differentiation (PPS threshold 20, transcript abundance threshold 10)

C. At the early pre-Tfh and GC-Tfh stages for pattern A of Tfh cell differentiation (PPS threshold 50, transcript abundance threshold 5)

D. At the early pre-Tfh and GC-Tfh stages for pattern B of Tfh cell differentiation (PPS threshold 50, transcript abundance threshold 10)

E. At the early pre-Tfh and GC-Tfh stages for pattern B of Tfh cell differentiation (PPS threshold 20, transcript abundance threshold 10)

F. At the early pre-Tfh and GC-Tfh stages for pattern B of Tfh cell differentiation (PPS threshold 50, transcript abundance threshold 5)

**Figure S3 – Extended functional overlap between Tfh cell subnetworks and known drivers of Tfh cell differentiation**

**Figure S4 - Most likely gene-level epigenomic patterns prior to network propagation for genes in Tfh subnetworks identified by network propagation**

A and B. Most likely gene-level ATAC-Seq pattern for genes in pattern B (A) and pattern A (B) Tfh subnetworks.

C and D. Most likely gene-level H3K4me1 pattern for genes in pattern B (C) and pattern A (D) Tfh subnetworks.

E and F. Most likely gene-level H3K4me3 pattern for genes in pattern B (E) and pattern A (F) Tfh subnetworks.

G and H. Most likely gene-level H3K27ac pattern for genes in pattern B (G) and pattern A (H) Tfh subnetworks.

**Figure S5 – Expression of genes in Tfh cell subnetworks in Tfh cells for patterns B and A respectively**

**Supplementary Data – Supplementary data accompanying the manuscript**

## Acknowledgements

Computational analyses were performed on the University of Pittsburgh Center for Research Computing (Pitt CRC) clusters and the Harvard Medical School Research Computing High Performance Cluster (HMS O2). This work was supported by a Pittsburgh Autoimmunity Center for Excellence in Research grant to J.D., NIH AI110495 and Ragon Institute Strategic Funding to S.P., by NIH AI113163 to V.M., and by the National Library of Medicine Biomedical Informatics and Data Science Research Training Program (T15LM007092) to V.V.

## References

Arunachalam, P.S., Wimmers, F., Mok, C.K.P., Perera, R.A.P.M., Scott, M., Hagan, T., Sigal, N., Feng, Y., Bristow, L., Tak-Yin Tsang, O., et al. (2020). Systems biological assessment of immunity to mild versus severe COVID-19 infection in humans. Science 369, 1210–1220.

Avsec, Ž., Agarwal, V., Visentin, D., Ledsam, J.R., Barwinska, A.G.-, Taylor, K.R., Assael, Y., Jumper, J., Kohli, P., and Kelley, D.R. Effective gene expression prediction from sequence by integrating long-range interactions.

Buenrostro, J.D., Giresi, P.G., Zaba, L.C., Chang, H.Y., and Greenleaf, W.J. (2013). Transposition of native chromatin for fast and sensitive epigenomic profiling of open chromatin, DNA-binding proteins and nucleosome position. Nat. Methods 10, 1213–1218.

Buenrostro, J.D., Wu, B., Chang, H.Y., and Greenleaf, W.J. (2015). ATAC-seq: A Method for Assaying Chromatin Accessibility Genome-Wide. Curr. Protoc. Mol. Biol. 21–29.

Chavele, K.-M., Merry, E., and Ehrenstein, M.R. (2015). Cutting edge: circulating plasmablasts induce the differentiation of human T follicular helper cells via IL-6 production. J. Immunol. 194, 2482–2485.

Cowen, L., Ideker, T., Raphael, B.J., and Sharan, R. (2017). Network propagation: a universal amplifier of genetic associations. Nat. Rev. Genet. 18, 551–562.

Crotty, S. (2014). T follicular helper cell differentiation, function, and roles in disease. Immunity 41, 529–542.

Cusick, M.E., Yu, H., Smolyar, A., Venkatesan, K., Carvunis, A.-R., Simonis, N., Rual, J.-F., Borick, H., Braun, P., Dreze, M., et al. (2009). Literature-curated protein interaction datasets. Nat. Methods 6, 39–46.

Das, J., and Yu, H. (2012). HINT: High-quality protein interactomes and their applications in understanding human disease. BMC Syst. Biol. 6, 92.

Das, J., Vo, T.V., Wei, X., Mellor, J.C., Tong, V., Degatano, A.G., Wang, X., Wang, L., Cordero, N.A., Kruer-Zerhusen, N., et al. (2013). Cross-species protein interactome mapping reveals species-specific wiring of stress response pathways. Sci. Signal. 6, ra38.

Das, J., Lee, H.R., Sagar, A., Fragoza, R., Liang, J., Wei, X., Wang, X., Mort, M., Stenson, P.D., Cooper, D.N., et al. (2014). Elucidating common structural features of human pathogenic variations using large-scale atomic-resolution protein networks. Hum. Mutat. 35, 585–593.

Elsner, R.A., and Shlomchik, M.J. (2019). IL-12 Blocks Tfh Cell Differentiation during Salmonella Infection, thereby Contributing to Germinal Center Suppression. Cell Rep. 29, 2796–2809.e5.

Filbin, M.R., Mehta, A., Schneider, A.M., Kays, K.R., Guess, J.R., Gentili, M., Fenyves, B.G., Charland, N.C., Gonye, A.L.K., Gushterova, I., et al. (2020). Plasma proteomics reveals tissuespecific cell death and mediators of cell-cell interactions in severe COVID-19 patients. bioRxiv.

Fragoza, R., Das, J., Wierbowski, S.D., Liang, J., Tran, T.N., Liang, S., Beltran, J.F., Rivera-Erick, C.A., Ye, K., Wang, T.-Y., et al. (2019). Extensive disruption of protein interactions by genetic variants across the allele frequency spectrum in human populations. Nat. Commun. 10, 4141.

Fulco, C.P., Munschauer, M., Anyoha, R., Munson, G., Grossman, S.R., Perez, E.M., Kane, M., Cleary, B., Lander, E.S., and Engreitz, J.M. (2016). Systematic mapping of functional enhancer-promoter connections with CRISPR interference. Science 354, 769–773.

Fulco, C.P., Nasser, J., Jones, T.R., Munson, G., Bergman, D.T., Subramanian, V., Grossman, S.R., Anyoha, R., Doughty, B.R., Patwardhan, T.A., et al. (2019). Activity-by-contact model of enhancer–promoter regulation from thousands of CRISPR perturbations. Nat. Genet. 51, 1664–1669.

Gordon, D.E., Jang, G.M., Bouhaddou, M., Xu, J., Obernier, K., White, K.M., O’Meara, M.J., Rezelj, V.V., Guo, J.Z., Swaney, D.L., et al. (2020). A SARS-CoV-2 protein interaction map reveals targets for drug repurposing. Nature 583, 459–468.

Hofree, M., Shen, J.P., Carter, H., Gross, A., and Ideker, T. (2013). Network-based stratification of tumor mutations. Nat. Methods 10, 1108–1115.

Horn, H., Lawrence, M.S., Chouinard, C.R., Shrestha, Y., Hu, J.X., Worstell, E., Shea, E., Ilic, N., Kim, E., Kamburov, A., et al. (2018). NetSig: network-based discovery from cancer genomes. Nat. Methods 15, 61–66.

Johnston, R.J., Poholek, A.C., DiToro, D., Yusuf, I., Eto, D., Barnett, B., Dent, A.L., Craft, J., and Crotty, S. (2009). Bcl6 and Blimp-1 are reciprocal and antagonistic regulators of T follicular helper cell differentiation. Science 325, 1006–1010.

Kaneko, N., Kuo, H.-H., Boucau, J., Farmer, J.R., Allard-Chamard, H., Mahajan, V.S., Piechocka-Trocha, A., Lefteri, K., Osborn, M., Bals, J., et al. (2020). Loss of Bcl-6-Expressing T Follicular Helper Cells and Germinal Centers in COVID-19. Cell 183, 143–157.e13.

Laing, A.G., Lorenc, A., Del Molino Del Barrio, I., Das, A., Fish, M., Monin, L., Muñoz-Ruiz, M., McKenzie, D.R., Hayday, T.S., Francos-Quijorna, I., et al. (2020). A dynamic COVID-19 immune signature includes associations with poor prognosis. Nat. Med. 26, 1623–1635.

Lei, X., Dong, X., Ma, R., Wang, W., Xiao, X., Tian, Z., Wang, C., Wang, Y., Li, L., Ren, L., et al. (2020). Activation and evasion of type I interferon responses by SARS-CoV-2. Nat. Commun. 11, 3810.

Leiserson, M.D.M., Eldridge, J.V., Ramachandran, S., and Raphael, B.J. (2013). Network analysis of GWAS data. Curr. Opin. Genet. Dev. 23, 602–610.

Leiserson, M.D.M., Vandin, F., Wu, H.-T., Dobson, J.R., Eldridge, J.V., Thomas, J.L., Papoutsaki, A., Kim, Y., Niu, B., McLellan, M., et al. (2015). Pan-cancer network analysis identifies combinations of rare somatic mutations across pathways and protein complexes. Nat. Genet. 47, 106–114.

Leng, N., Dawson, J.A., Thomson, J.A., Ruotti, V., Rissman, A.I., Smits, B.M.G., Haag, J.D., Gould, M.N., Stewart, R.M., and Kendziorski, C. (2013). EBSeq: an empirical Bayes hierarchical model for inference in RNA-seq experiments. Bioinformatics 29, 1035–1043.

Marasco, M., Berteotti, A., Weyershaeuser, J., Thorausch, N., Sikorska, J., Krausze, J., Brandt, H.J., Kirkpatrick, J., Rios, P., Schamel, W.W., et al. (2020). Molecular mechanism of SHP2 activation by PD-1 stimulation. Sci Adv 6, eaay4458.

Nasser, J., Bergman, D.T., Fulco, C.P., Guckelberger, P., Doughty, B.R., Patwardhan, T.A., Jones, T.R., Nguyen, T.H., Ulirsch, J.C., Lekschas, F., et al. (2021). Genome-wide enhancer maps link risk variants to disease genes. Nature.

Overmyer, K.A., Shishkova, E., Miller, I.J., Balnis, J., Bernstein, M.N., Peters-Clarke, T.M., Meyer, J.G., Quan, Q., Muehlbauer, L.K., Trujillo, E.A., et al. (2021). Large-Scale Multi-omic Analysis of COVID-19 Severity. Cell Syst 12, 23–40.e7.

Reyna, M.A., Leiserson, M.D.M., and Raphael, B.J. (2018). Hierarchical HotNet: identifying hierarchies of altered subnetworks. Bioinformatics 34, i972–i980.

Schmitt, N., Morita, R., Bourdery, L., Bentebibel, S.E., Zurawski, S.M., Banchereau, J., and Ueno, H. (2009). Human dendritic cells induce the differentiation of interleukin-21-producing T follicular helper-like cells through interleukin-12. Immunity 31, 158–169.

Schmitt, N., Liu, Y., Bentebibel, S.-E., Munagala, I., Bourdery, L., Venuprasad, K., Banchereau, J., and Ueno, H. (2014). The cytokine TGF-β co-opts signaling via STAT3-STAT4 to promote the differentiation of human TFH cells. Nat. Immunol. 15, 856–865.

Schrock, D.C., Leddon, S.A., Hughson, A., Miller, J., Lacy-Hulbert, A., and Fowell, D.J. (2019). Pivotal role for αV integrins in sustained Tfh support of the germinal center response for long-lived plasma cell generation. Proc. Natl. Acad. Sci. U. S. A. 116, 4462–4470.

Skene, P.J., and Henikoff, S. (2017). An efficient targeted nuclease strategy for high-resolution mapping of DNA binding sites. Elife 6, e21856.

Subramanian, A., Tamayo, P., Mootha, V.K., Mukherjee, S., Ebert, B.L., Gillette, M.A., Paulovich, A., Pomeroy, S.L., Golub, T.R., Lander, E.S., et al. (2005). Gene set enrichment analysis: a knowledge-based approach for interpreting genome-wide expression profiles. Proc. Natl. Acad. Sci. U. S. A. 102, 15545–15550.

Vandin, F., Upfal, E., and Raphael, B.J. (2011). Algorithms for detecting significantly mutated pathways in cancer. J. Comput. Biol. 18, 507–522.

Vidal, M., Cusick, M.E., and Barabási, A.-L. (2011). Interactome networks and human disease. Cell 144, 986–998.

Vinuesa, C.G., Linterman, M.A., Yu, D., and MacLennan, I.C.M. (2016). Follicular Helper T Cells. Annu. Rev. Immunol. 34, 335–368.

Vo, T.V., Das, J., Meyer, M.J., Cordero, N.A., Akturk, N., Wei, X., Fair, B.J., Degatano, A.G., Fragoza, R., Liu, L.G., et al. (2016). A Proteome-wide Fission Yeast Interactome Reveals Network Evolution Principles from Yeasts to Human. Cell 164, 310–323.

Wang, X., Wei, X., Thijssen, B., Das, J., Lipkin, S.M., and Yu, H. (2012). Three-dimensional reconstruction of protein networks provides insight into human genetic disease. Nat. Biotechnol. 30, 159–164.

Wei, X., Das, J., Fragoza, R., Liang, J., Bastos de Oliveira, F.M., Lee, H.R., Wang, X., Mort, M., Stenson, P.D., Cooper, D.N., et al. (2014). A massively parallel pipeline to clone DNA variants and examine molecular phenotypes of human disease mutations. PLoS Genet. 10, e1004819.

Whalen, S., Truty, R.M., and Pollard, K.S. (2016). Enhancer-promoter interactions are encoded by complex genomic signatures on looping chromatin. Nat. Genet. 48, 488–496.

Yi, S., Lin, S., Li, Y., Zhao, W., Mills, G.B., and Sahni, N. (2017). Functional variomics and network perturbation: connecting genotype to phenotype in cancer. Nat. Rev. Genet. 18, 395–410.

Yu, B., Zhang, K., Milner, J.J., Toma, C., Chen, R., Scott-Browne, J.P., Pereira, R.M., Crotty, S., Chang, J.T., Pipkin, M.E., et al. (2017). Epigenetic landscapes reveal transcription factors that regulate CD8+ T cell differentiation. Nat. Immunol. 18, 573–582.

Zuin, J., Roth, G., Zhan, Y., Cramard, J., Redolfi, J., Piskadlo, E., Mach, P., Kryzhanovska, M., Tihanyi, G., Kohler, H., et al. (2021). Nonlinear control of transcription through enhancerpromoter interactions.

