## Supplementary Figures for "Network-based integration of epigenetic landscapes unveils molecular programs underlying human T follicular helper cell differentiation"

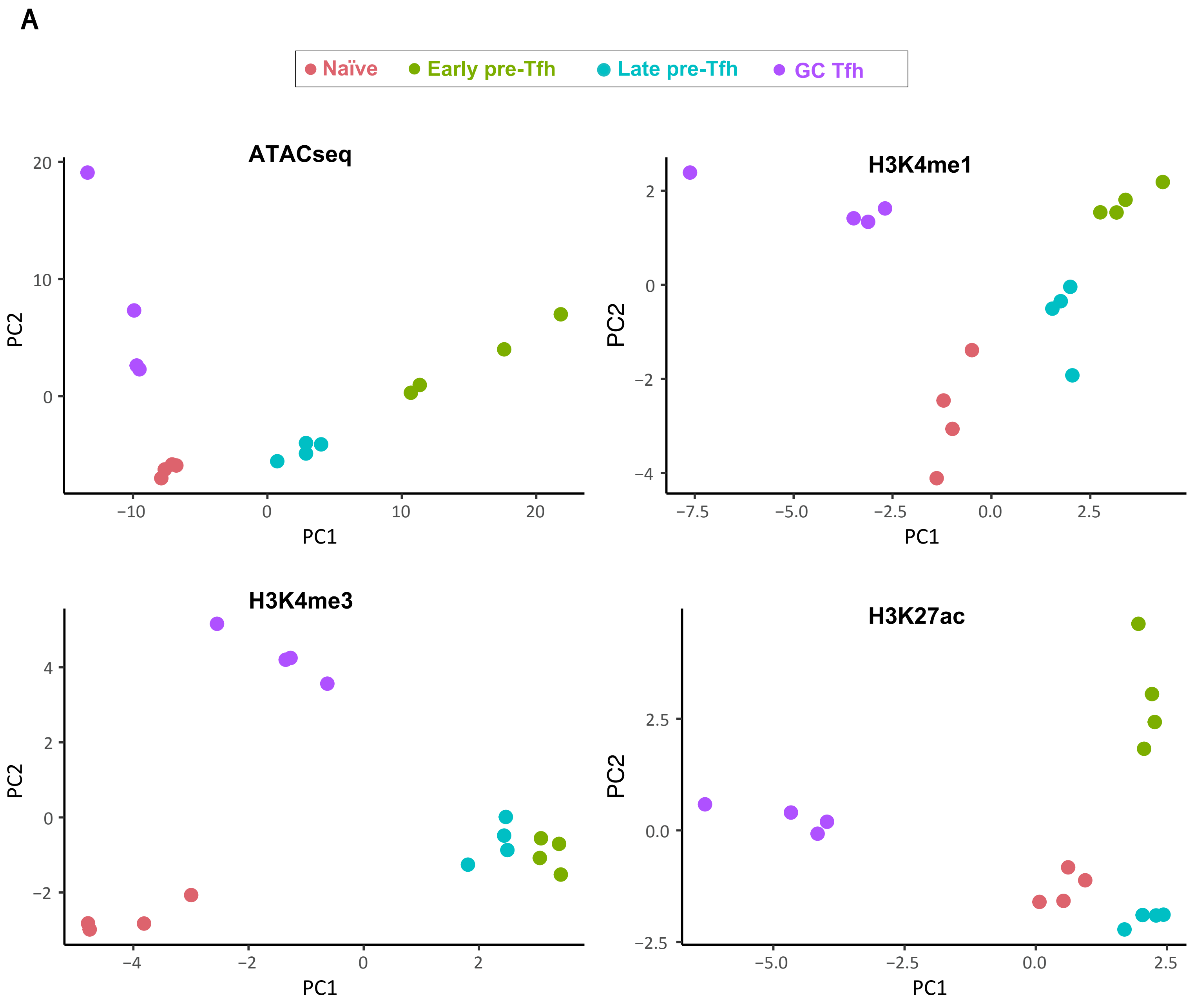
A
Naïve
Early pre-Tfh
Late pre-Tfh
GC Tfh
ATACseq
H3K4me1
H3K4me3
H3K27ac
PC1
PC2

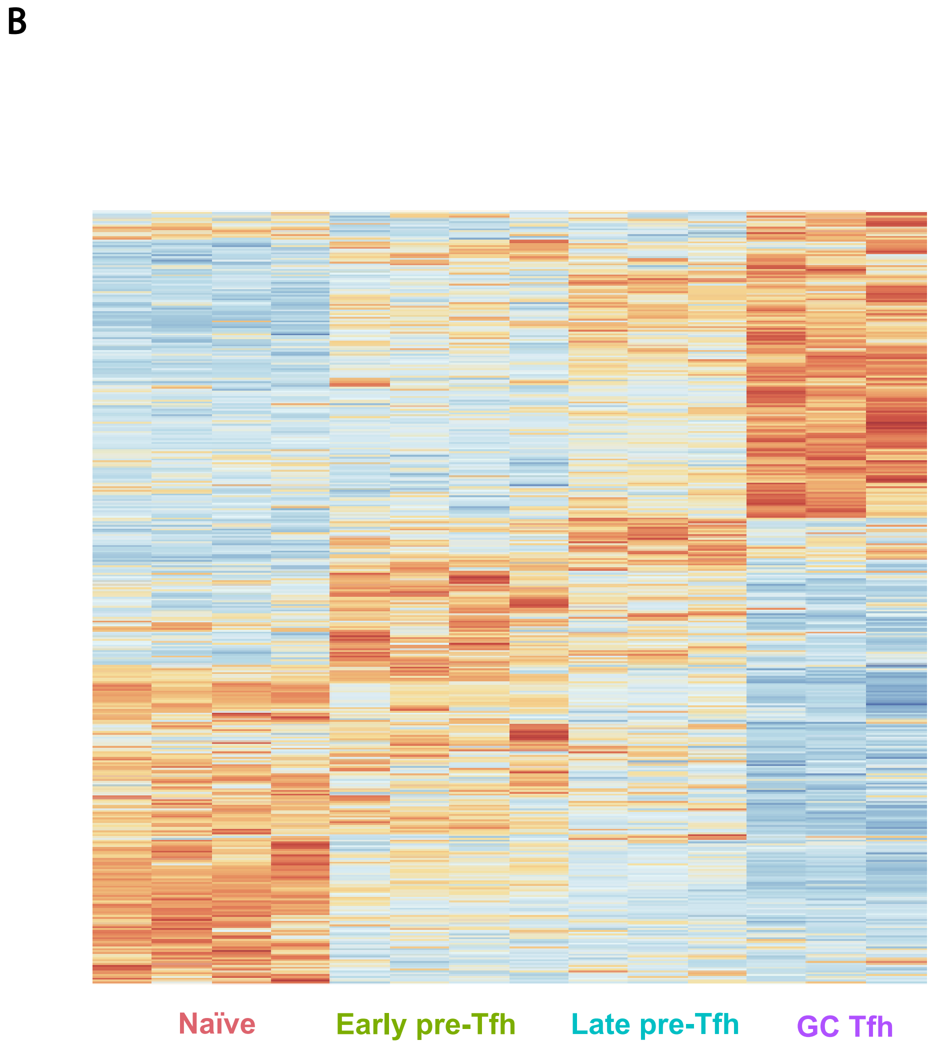
B
Naïve
Early pre-Tfh
Late pre-Tfh
GC Tfh

## A

**early pre-Tfh** — **Pattern A**

Gene Scores

| Transcript abundance: p = 0.0356 | Low | High |
|---|---|---|
| Low | 4941 | 385 |
| High | 2101 | 194 |

**GC Tfh** — **Pattern A**

Gene Score

| Transcript abundance: p = 0.0057 | Low | High |
|---|---|---|
| Low | 5084 | 389 |
| High | 1958 | 190 |

## B

Gene Scores

| Transcript abundance: p = 0.003 | Low | High |
|---|---|---|
| Low | 4447 | 879 |
| High | 1856 | 439 |

Gene Score

| Transcript abundance: p = 0.0073 | Low | High |
|---|---|---|
| Low | 4567 | 906 |
| High | 1736 | 412 |

## C

Gene Scores

| Transcript abundance: p = 0.0095 | Low | High |
|---|---|---|
| Low | 4379 | 331 |
| High | 2663 | 248 |

Gene Score

| Transcript abundance: p = 0.0203 | Low | High |
|---|---|---|
| Low | 4455 | 341 |
| High | 2587 | 238 |

## D

**early pre-Tfh** — **Pattern B**

Gene Score

| Transcript abundance: p=0.0001 | Low | High |
|---|---|---|
| Low | 8028 | 1044 |
| High | 3139 | 577 |

**GC Tfh** — **Pattern B**

Gene Score

| Transcript abundance: p=0.0001 | Low | High |
|---|---|---|
| Low | 8237 | 1079 |
| High | 2930 | 542 |

## E

Gene Score

| Transcript abundance: p=0.0001 | Low | High |
|---|---|---|
| Low | 6537 | 2535 |
| High | 2439 | 1277 |

Gene Score

| Transcript abundance: p=0.0001 | Low | High |
|---|---|---|
| Low | 6682 | 2634 |
| High | 2294 | 1178 |

## F

Gene Score

| Transcript abundance: p=0.0001 | Low | High |
|---|---|---|
| Low | 7145 | 877 |
| High | 4022 | 744 |

Gene Score

| Transcript abundance: p=0.0001 | Low | High |
|---|---|---|
| Low | 7297 | 915 |
| High | 3870 | 706 |

A

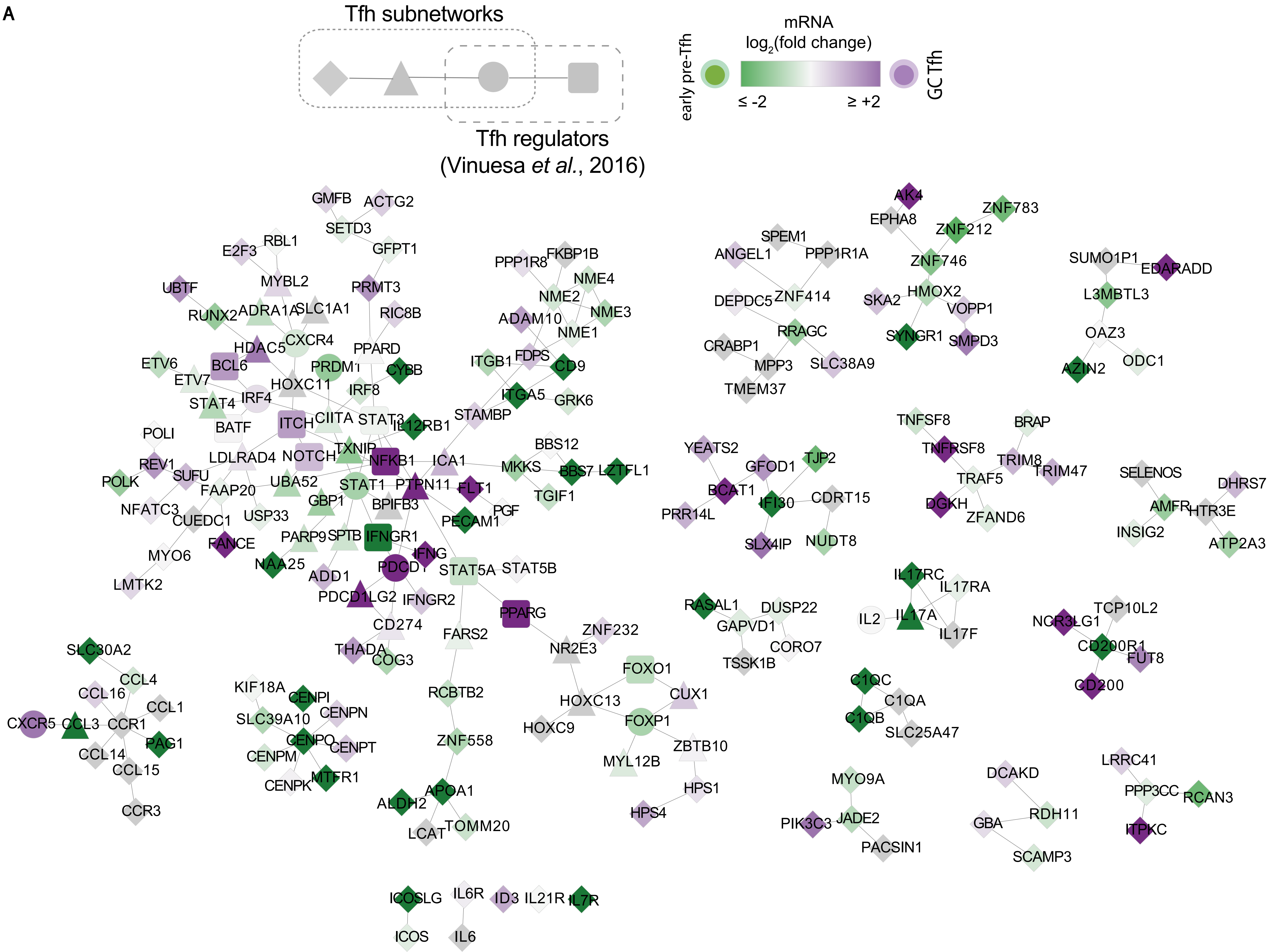

Tfh subnetworks
Tfh regulators
(Vinuesa et al., 2016)
mRNA
log2(fold change)
early pre-Tfh
GC Tfh
≤ -2
≥ +2
GMFB ACTG2 SETD3 GFPT1 RBL1 E2F3 MYBL2 UBTF PRMT3 RUNX2 ADRA1A SLC1A1 RIC8B HDAC5 CXCR4 PPARD ETV6 BCL6 PRDM1 CYBB ETV7 HOXC11 IRF8 STAT4 IRF4 ITCH CIITA STAT3 IL12RB1 BATF POLI LDLRAD4 NOTCH TXNIP NFKB1 ICA1 REV1 SUFU POLK FAAP20 UBA52 STAT1 PTPN11 FLT1 GBP1 BPIFB3 NFATC3 CUEDC1 USP33 PGF PECAM1 FANCE PARP9 SPTB IFNGR1 MYO6 IFNG NAA25 PDCD1 STAT5A STAT5B ADD1 LMTK2 PDCD1LG2 IFNGR2 PPARG CD274 ZNF232 FARS2 THADA NR2E3 COG3
PPP1R8 FKBP1B NME4 NME2 NME3 ADAM10 NME1 ITGB1 FDPS CD9 ITGA5 GRK6 STAMBP BBS12 MKKS BBS7 LZTFL1 TGIF1
SPEM1 ANGEL1 PPP1R1A DEPDC5 ZNF414 RRAGC CRABP1 MPP3 SLC38A9 TMEM37
AK4 EPHA8 ZNF783 ZNF212 ZNF746 HMOX2 SKA2 VOPP1 SYNGR1 SMPD3
SUMO1P1 EDARADD L3MBTL3 OAZ3 AZIN2 ODC1
YEATS2 GFOD1 TJP2 BCAT1 IFI30 CDRT15 PRR14L SLX4IP NUDT8
TNFSF8 BRAP TNFRSF8 TRIM8 TRIM47 TRAF5 DGKH ZFAND6
SELENOS DHRS7 AMFR HTR3E INSIG2 ATP2A3
IL17RC IL17RA IL2 IL17A IL17F
RASAL1 DUSP22 GAPVD1 CORO7 TSSK1B
TCP10L2 NCR3LG1 CD200R1 FUT8 CD200
SLC30A2 CCL4 CCL16 CCL1 CXCR5 CCL3 CCR1 PAG1 CCL14 CCL15 CCR3
KIF18A CENPI CENPN SLC39A10 CENPO CENPT CENPM CENPK MTFR1
RCBTB2 ZNF558 APOA1 ALDH2 LCAT TOMM20
FOXO1 CUX1 HOXC13 HOXC9 FOXP1 ZBTB10 MYL12B HPS1 HPS4
C1QC C1QA C1QB SLC25A47
MYO9A PIK3C3 JADE2 PACSIN1
DCAKD RDH11 GBA SCAMP3
LRRC41 PPP3CC RCAN3 ITPKC
ICOSLG IL6R ID3 IL21R IL7R ICOS IL6

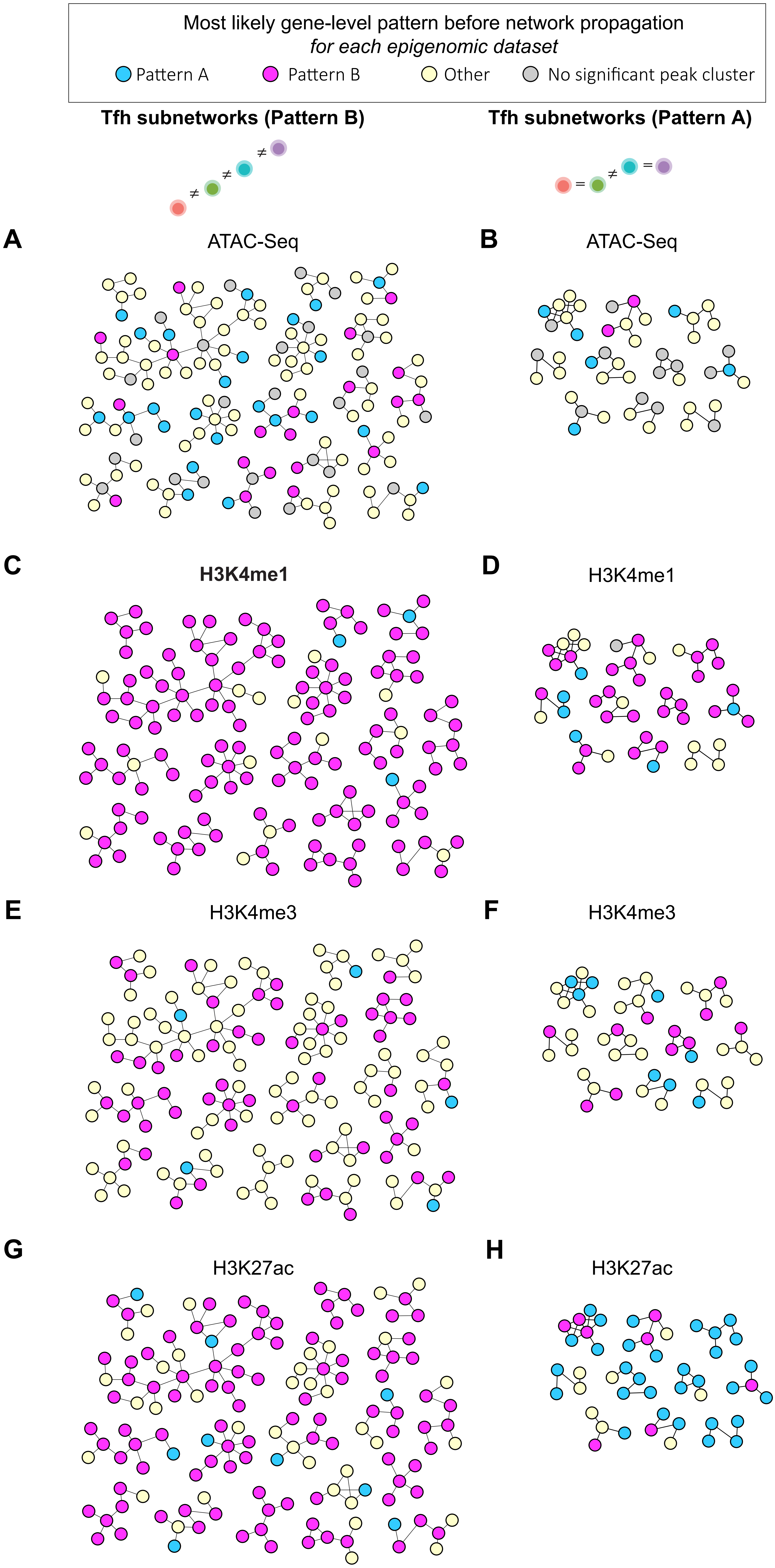

Most likely gene-level pattern before network propagation
for each epigenomic dataset
Pattern A
Pattern B
Other
No significant peak cluster
Tfh subnetworks (Pattern B)
Tfh subnetworks (Pattern A)
≠
≠
≠
=
≠
=
A
ATAC-Seq
B
ATAC-Seq
C
H3K4me1
D
H3K4me1
E
H3K4me3
F
H3K4me3
G
H3K27ac
H
H3K27ac

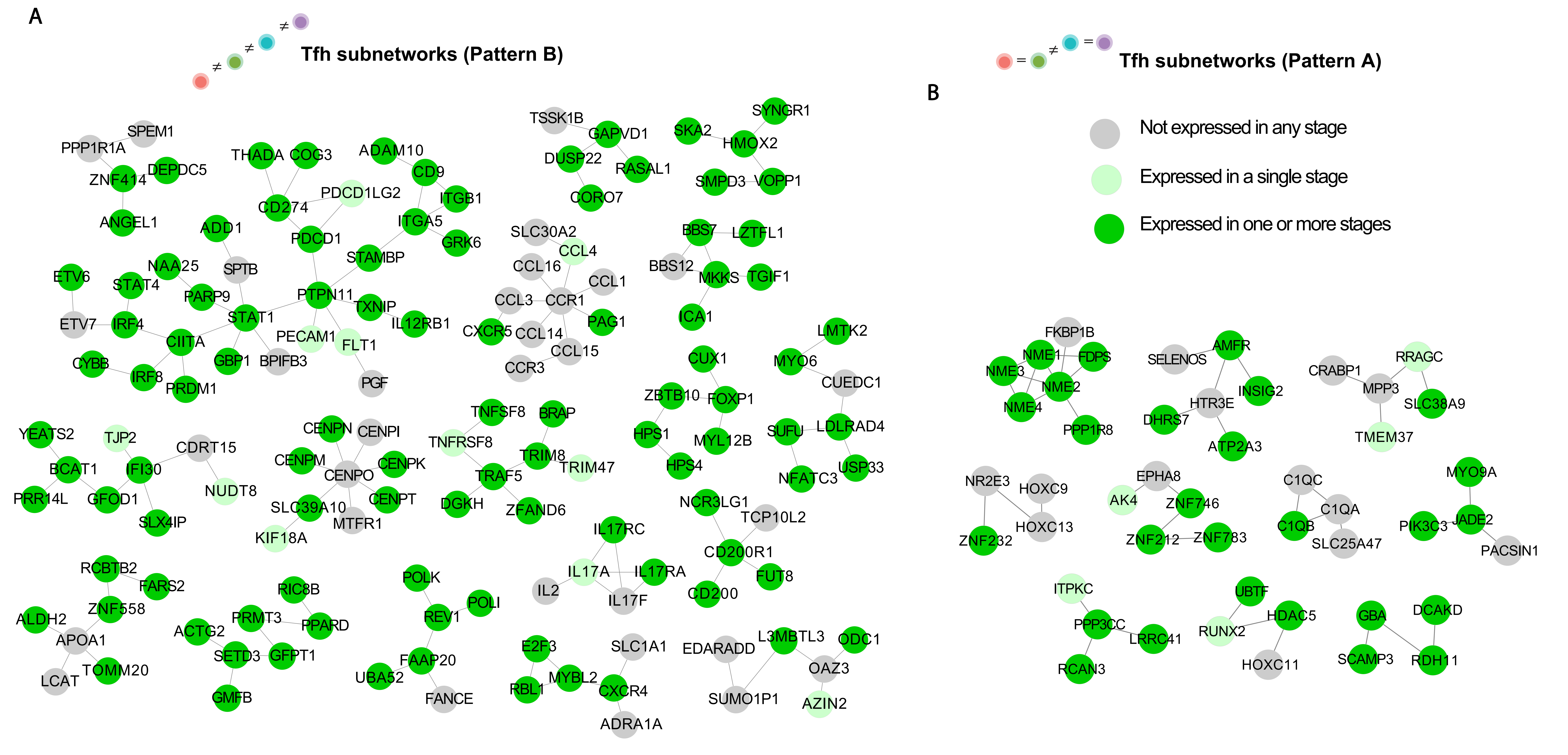
A
Tfh subnetworks (Pattern B)
Tfh subnetworks (Pattern A)
B
Not expressed in any stage
Expressed in a single stage
Expressed in one or more stages
SPEM1
PPP1R1A
DEPDC5
ZNF414
ANGEL1
THADA
COG3
ADAM10
CD9
ITGB1
PDCD1LG2
CD274
PDCD1
ITGA5
GRK6
STAMBP
ADD1
SPTB
NAA25
ETV6
STAT4
PARP9
PTPN11
TXNIP
IL12RB1
ETV7
IRF4
STAT1
CIITA
PECAM1
FLT1
PGF
CYBB
IRF8
PRDM1
GBP1
BPIFB3
TSSK1B
GAPVD1
DUSP22
RASAL1
CORO7
SKA2
SYNGR1
HMOX2
SMPD3
VOPP1
SLC30A2
CCL4
CCL16
CCL1
CCL3
CCR1
PAG1
CXCR5
CCL14
CCL15
CCR3
BBS7
LZTFL1
BBS12
MKKS
TGIF1
ICA1
CUX1
LMTK2
MYO6
CUEDC1
ZBTB10
FOXP1
HPS1
MYL12B
HPS4
SUFU
LDLRAD4
NFATC3
USP33
YEATS2
TJP2
CDRT15
BCAT1
IFI30
NUDT8
PRR14L
GFOD1
SLX4IP
CENPN
CENPI
CENPM
CENPO
CENPK
SLC39A10
CENPT
MTFR1
KIF18A
TNFSF8
BRAP
TNFRSF8
TRIM8
TRIM47
TRAF5
DGKH
ZFAND6
NCR3LG1
TCP10L2
IL17RC
CD200R1
IL17A
IL17RA
IL2
IL17F
FUT8
CD200
RCBTB2
FARS2
ZNF558
ALDH2
APOA1
LCAT
TOMM20
RIC8B
PRMT3
PPARD
ACTG2
SETD3
GFPT1
GMFB
POLK
POLI
REV1
FAAP20
UBA52
FANCE
E2F3
SLC1A1
MYBL2
RBL1
CXCR4
ADRA1A
EDARADD
L3MBTL3
ODC1
OAZ3
SUMO1P1
AZIN2
FKBP1B
NME1
FDPS
NME3
NME2
NME4
PPP1R8
SELENOS
AMFR
INSIG2
HTR3E
DHRS7
ATP2A3
CRABP1
RRAGC
MPP3
SLC38A9
TMEM37
NR2E3
HOXC9
HOXC13
ZNF232
EPHA8
AK4
ZNF746
ZNF212
ZNF783
C1QC
C1QA
C1QB
SLC25A47
MYO9A
PIK3C3
JADE2
PACSIN1
ITPKC
PPP3CC
LRRC41
RCAN3
UBTF
RUNX2
HDAC5
HOXC11
GBA
DCAKD
SCAMP3
RDH11
